# Antibiotic condensates activate magnesium signaling to drive drug tolerance

**DOI:** 10.64898/2026.08.17.745126

**Authors:** Yuefeng Ma, Wen Yu, Eunchae Moon, Shanlong Li, Zongru Li, Yanrun Zhou, James Wang, Jiaxin Chen, Xinrui Liu, Nicholas Wilson, Erica Lantelme, Ellie Berkland, Jianhan Chen, Gürol M. Süel, Yifan Dai

**Affiliations:** Department of Biomedical Engineering, Washington University in St. Louis, Saint Louis, MO, 63130; Department of Molecular Biology, School of Biological Sciences, University of California San Diego, La Jolla, CA, 92121; Department of Chemistry, University of Massachusetts, Amherst, MA, 01003; Department of Chemistry, Washington University in St. Louis, Saint Louis, MO, 63130; Center for Biomolecular Condensates, Washington University in St. Louis, Saint Louis, MO, 63130; Department of Pathology and Immunology, Washington University in St. Louis, Saint Louis, MO, 63110, USA

## Abstract

The spatial distribution of small molecules within cells shapes their biological activity, yet these distributions are generally assumed to be governed passively by reaction-driven electrochemical gradients. Here we show that aminoglycoside antibiotics actively control their own subcellular organization by undergoing phase separation with RNAs. Combining in vitro reconstitution, bacterial assays, and molecular dynamics simulations, we discovered that aminoglycosides coacervate with RNA through multivalent electrostatic interactions, displacing and releasing RNA-bound Mg^2+^. This condensate-dependent Mg^2+^ release remodels the cytosolic labile Mg^2+^ pool and activates magnesium signaling. This effect dampens the magnesium-starvation regulation, sustains ribosome activity, and shifts the cellular electrochemical state, promoting bacterial fitness. Because condensation occurs only above a defined concentration threshold, it generates a non-monotonic dose-response in which higher antibiotic concentrations paradoxically enhance bacterial survival. This antibiotic condensate-dependent Mg^2+^ signaling confers tolerance to multiple ribosome-targeting antibiotics simultaneously even in cells lacking resistance gene, while condensate dissolution restores antibiotic efficacy. Our findings establish antibiotic-driven phase separation as a previously unrecognized mechanism to encode cellular signaling and identify antibiotic condensates as a distinct functional unit underlying drug tolerance.

## Main Text

Understanding cellular organization principles is reshaping our knowledge of how biological systems function^1^. Over the past decade, studies have established that dynamic and multivalent biomacromolecules, particularly intrinsically disordered proteins and RNAs^2–5^, serve as the key drivers modulating the spatiotemporal organization of cellular processes, from transcription to stress responses^6–8^. Through phase transitions^9,10^, these biomacromolecules scaffold the formation of biomolecular condensates and recruit specific biomolecules into the condensates, encoding selective cellular functions^11,12^.

Yet cellular organization is not limited to macromolecules. Small molecules, such as ions, metabolites, also exhibit distinct spatiotemporal distribution within cells^13–16^, and their distributions, which shape local physicochemical environments, regulate biological activities dynamically^17–20^. However, studies have largely assumed that the distribution of small molecules is passively governed by reaction-driven concentration gradients^21^, which arise from electrochemical potential differences determined by their charge and solvation properties, as exemplified by the chemiosmotic theory^22,23^. Whether small molecules can actively control their own distribution, the mechanisms governing this process and the cellular functions of their self-organization remain largely explored.

This question is particularly important for antibiotics, the resistance and tolerance of which is a global health emergency^24,25^. The biological activity of antibiotics depends not only on their chemical structure but also on where they accumulate within cells because most antibiotic targets are localized to distinct subcellular regions^26–28^. Thus, uncovering the principles that govern small molecule organization in space and time may provide new insights into the mechanisms of antibiotic action and tolerance.

Here we demonstrate that aminoglycosides, a key category of bactericidal antibiotics that possess a central aminocyclitol linked to multiple amino sugars by pseudoglycosidic bonds^29^, can regulate their own spatiotemporal distributions in cells through co-condensing with RNAs. This phase transition process releases the bound Mg^2+^ ions from the RNAs and modulates the cytosolic Mg^2+^ distribution, which is studied through in vitro reconstitution assays, cellular assays and molecular dynamics simulations. The condensate-dependent Mg^2+^ release activates multiple cellular pathways through Mg^2+^ signaling as examined with RNA-sequencing, enhancing ribosome activity in a phase separation-dependent manner. This explains the paradoxical antibiotic response^30^, whereby survival is enhanced at higher drug concentrations, producing a non-monotonic relationship between gentamicin dose and cell viability. Dissolving the antibiotic condensates successfully restored antibiotic efficacy. Our study establishes that antibiotics can drive phase separation, a previously unrecognized functioning mechanism, to redistribute labile Mg^2+^ ions and activate downstream electrochemical signaling to sustain ribosome activities under antibiotic stress, thus mediating multi-drug tolerance in cells without any antibiotic resistance genes. This finding enables prediction of the survival window from the phase-separation threshold of aminoglycosides, which reveals a new gene-independent mechanism underlying broad drug tolerance and provides condensate disruption as a strategy for restoring antibiotic efficacy.

## Results

### Aminoglycosides drive phase separation with RNA

A key molecular principle underlying the phase transition of biomacromolecules is multivalency^9,31^ (**Fig. 1a**). In biomacromolecules, such as intrinsically disordered proteins^32^, multiple different “sticker”-type of residues (n>2), such as aromatic residues, can engage in multiple intra/inter-chain interactions^33,34^, thereby driving phase transition and condensate formation. The same principle should be also applicable to the phase separation involving small molecules^35,36^, such as the widely used adenosine triphosphate for complex coacervations with polyamines.

**Figure 1.**
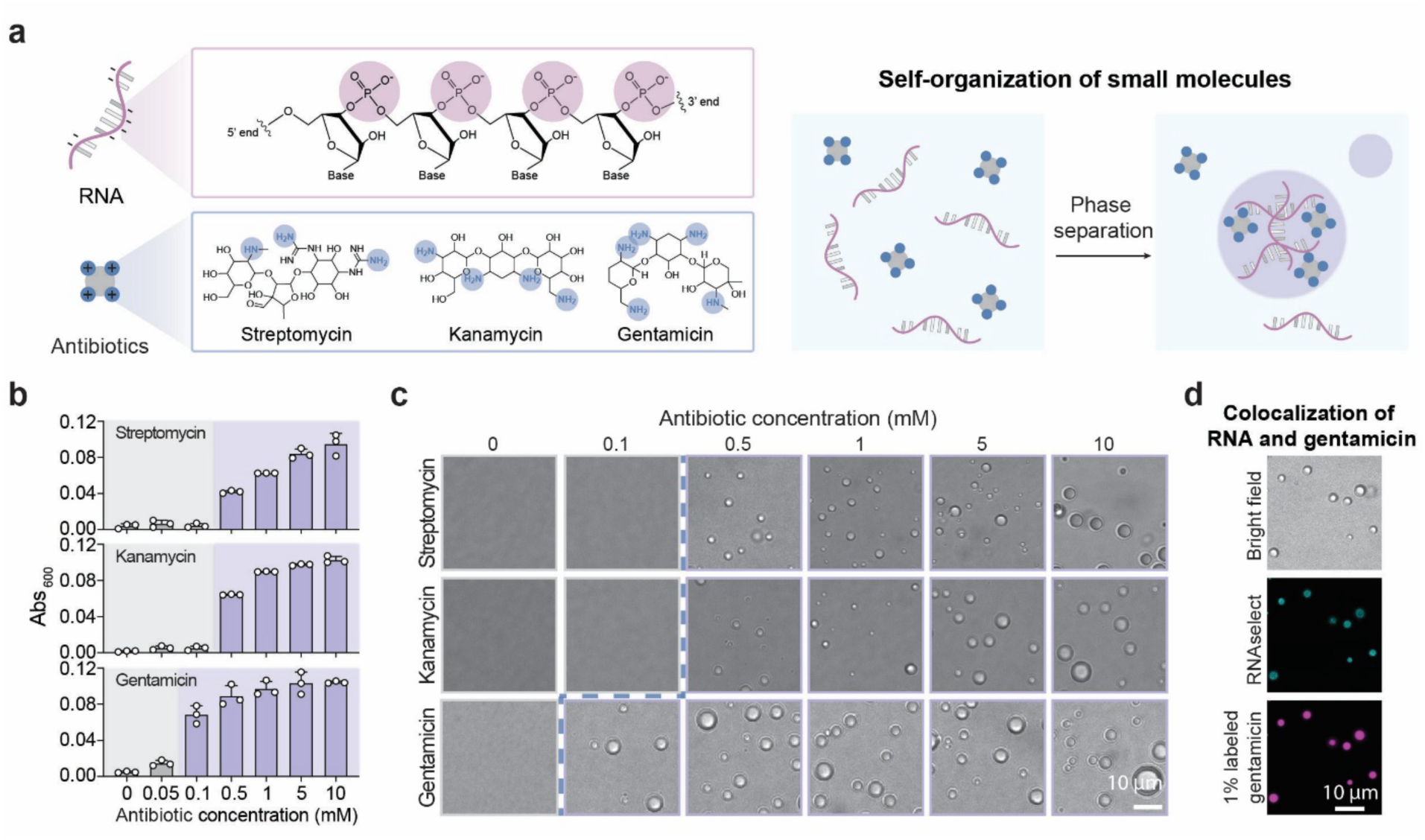
Aminoglycosides drive biomolecular condensate formation through RNA condensation. **a,** Chemical structures of RNA and the aminoglycoside antibiotics streptomycin, kanamycin and gentamicin, with a schematic of their condensation. Charge complementarity between polyanionic RNA and polycationic antibiotics enables multivalent electrostatic interactions, promoting biomolecular condensate formation. **b,** Absorbance measurements at 600 nm indicate *in vitro* biomolecular condensate formation. Grey and purple shading denote conditions without and with condensate formation, respectively. **c,** Confocal images showing *in vitro* biomolecular condensate formation across different concentrations of antibiotics. Confocal images with grey and purple border denote conditions without and with condensate formation, respectively. Dash lines delineate the phase-separation boundaries for streptomycin, kanamycin and gentamicin. Scale bar represents 10 μm. **d,** Confocal fluorescence images of gentamicin–RNA condensates showing the colocalization of gentamicin and RNA *in vitro*. Gentamicin and RNA are visualized using 1% (w/w) Texas Red-labelled gentamicin and 0.1 μM RNAselect, respectively. Scale bar represents 10 μm.

We therefore asked whether aminoglycosides, positively charged, multi-amine antibiotics built on 4,5- or 4,6-disubstituted deoxystreptamine scaffolds, could similarly undergo phase separation with negatively charged RNAs. To this end, we tested the abilities of the widely used gentamicin, kanamycin and streptomycin on mediating phase transition with a model polyA RNA by evaluating the changes of solution turbidity in a concentration-dependent manner (**Fig. 1b**). Under the same buffer condition and a fixed RNA concentration, with confocal microscopy using phase contrast, we observed antibiotic-concentration-dependent condensate formation (**Fig. 1c**), consistent with the threshold concentration identified by turbidity measurements. To evaluate whether the phase transition behavior depends on the number of charges on the aminoglycosides, we compared the phase diagram of kanamycin with that of gentamicin^37,38^, which is less protonated than gentamicin at physiological condition, in the presence of the same RNA concentration, and found that the threshold concentration increased substantially (**Fig. 1c**). Similarly, under the same aminoglycosides concentration, the phase behavior is modulated by the RNA concentration in the system (**Supplementary Fig. 1a**). This observation confirms that antibiotic condensates can be formed through complex coacervation^39^, governed by the charge valency and concentration of the antibiotics and RNA.

To further confirm the molecular components mediating condensation, we doped a fluorescence-labeled gentamicin analog, iFluor594 gentamicin conjugate (GTIF594)^40^, and a fluorescent RNA dye, RNASelect^41^, into the gentamicin-RNA mixture (**Fig. 1d**). With confocal microscopy, we observed the colocalization between RNA and gentamicin analog in the condensates. Lastly, we examined the capability of gentamicin to condense DNA or polyphosphates^42^, which are key negatively charged components in bacteria. However, unlike RNA, which condenses at a low concentration, physiologically relevant concentrations of DNA or polyphosphate failed to drive condensation at the same gentamicin level (**Supplementary Fig. 1b and 1c**). Further, when we mixed the RNA, DNA and polyphosphate together with gentamicin at their physiologically relevant concentration, we observed condensate formation, and confirmed the key roles of RNA and gentamicin on driving phase separation with the fluorescence assay (**Supplementary Fig. 1d**). This assay suggests that RNA can favorably interact with gentamicin, possibly due to its conformational flexibility and its intrinsic capability to form inter/intra-RNA interactions facilitated by condensation^43,44^. Together, these results establish that aminoglycosides can self-organize by favorably driving phase separation with RNA.

### Phase transition modulates labile ion availability

In an electrolyte solution, the transition from a homogeneous solution to a two-phase aqueous system shifts ion distribution^39,45–48^. Before the coacervation process between aminoglycosides and RNA, the RNA should be electrostatically screened by small cations, so that the interaction between amino groups on the antibiotics and the phosphate backbones of RNA might result in the release of small cations from the RNA (**Fig. 2a**). The specific cation that is critical to RNA functions in bacteria is Mg^2+^ ^49,50^, which is also a key signaling ion that regulates diverse bacterial functions^50,51^. Thus, we hypothesized that aminoglycosides induced RNA phase transition might modulate ion signaling.

**Figure 2.**
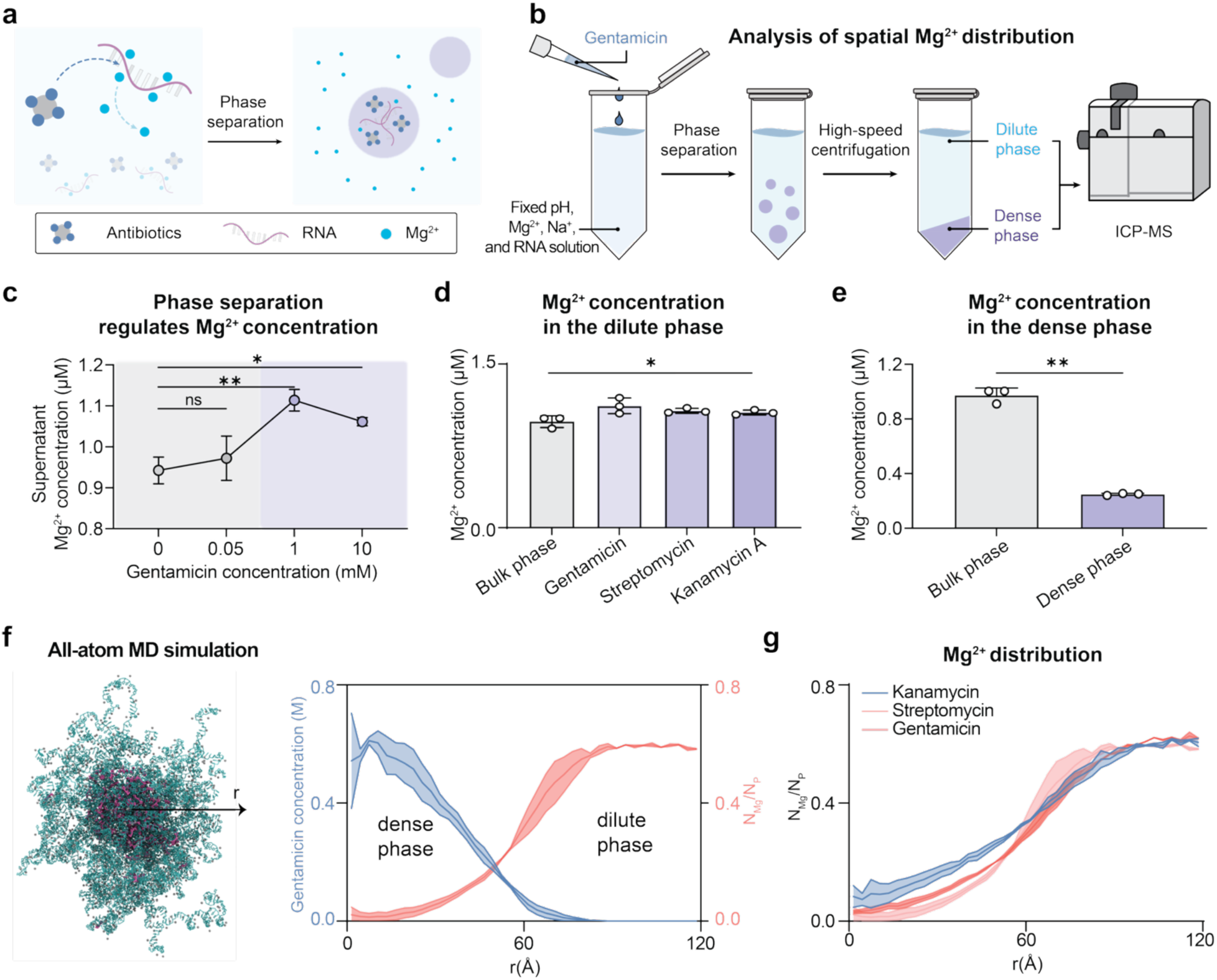
Aminoglycoside-RNA condensate formation regulates spatial Mg^2+^ distribution. **a,** Schematic of the formation of antibiotic–RNA condensates, leading to the release of magnesium ions from the RNA template. Due to their higher affinity for the RNA backbone, antibiotics competitively displace magnesium ions based on condensate formation. **b,** Schematic workflow for *in vitro* measurement of magnesium concentrations by ICP–MS. Gentamicin is added to a buffer containing fixed concentrations of Mg^2+^, Na⁺ and RNA at a defined pH. After incubation for 1 h at room temperature, samples are centrifuged to separate the dilute and dense phases. The supernatant (dilute phase) and pellet (dense phase) are collected separately for ion concentration measurements. **c,** Evaluation of supernatant magnesium concentrations following the addition of different concentrations of gentamicin. Grey and purple shading denote conditions without and with condensate formation, respectively. Data is represented as mean ± SD. N = 3 biological replicates. Two-tailed t test for statistical analysis. **d,** Evaluation of supernatant magnesium concentrations following the addition of 1 mM gentamicin, streptomycin, or kanamycin A. For the bulk-phase control, an equivalent volume of antibiotic-free buffer is added. N = 3 biological replicates. Statistical significance was determined using an ordinary one-way ANOVA test. **e,** Comparison of magnesium concentrations in the dense phase of gentamicin–RNA condensates and in the bulk phase. For the gentamicin-RNA condensate formation, gentamicin is added at a final concentration of 1 mM into a buffer containing fixed concentrations of Mg^2+^, Na⁺ and RNA at a defined pH. For the bulk-phase control, an equivalent volume of antibiotic-free buffer is added. N = 3 biological replicates. Two-tailed t test for statistical analysis. **f,** Representative snapshot of rA200-kanamycin-Mg cluster. The arrow indicates the distance (r) from the center of the condensate to the measurement position. Radial distributions of kanamycin concentration (purple) and Mg number per phosphate (red). **g,** Effects of streptomycin (red), kanamycin (purple), and gentamicin (green) on the number of Mg within condensates. Error bars (shadows) were estimated from block deviations. The exclusive effect is enhanced with the increase of net charges of the antibiotics.

To assess this, we prepared solutions containing the same concentrations of Mg^2+^ and RNA but varying concentrations of gentamicin, and quantified the accessible Mg^2+^ in the supernatant after sedimentation using inductively coupled plasma mass spectrometry (ICP–MS) ^48,52^ (**Fig. 2b**). Before surpassing the saturation threshold for phase transition, sedimentation assay does not produce bulk phase separation, and the measured Mg^2+^ therefore reflects the total Mg^2+^ present in solution. After passing the phase transition boundary, however, sedimentation induces bulk phase separation, such that if there is a change in Mg^2+^ detected in the supernatant, this difference is the concentration gradient generates by phase transition, which corresponds to the released labile Mg^2+^.

We found that before phase transition (C_gentamicin_ = 0 to 0.05 mM), increasing gentamicin concentration in the solution did not alter the amount of Mg^2+^ probed at a fixed RNA concentration (**Fig. 2c**), which suggests that Mg^2+^ is tightly regulated by the electrostatic interactions with RNAs in the solvated state. However, once the gentamicin concentration surpasses the phase transition threshold (C_gentamicin_ >0.1 mM), the amount of Mg^2+^ detected noticeably increased. Further, we identified that at the same aminoglycoside concentration that crosses the phase boundary, the apparent charge of the aminoglycoside modulates the amount of Mg^2+^ released (**Fig. 2d**).

Since the total volume fraction of condensates in the solution is typically limited (∼0.01%)^53^, which is not comparable to the volume fraction of condensates in bacteria (∼10-20%)^48^, to further verify this result, we next analyzed the Mg^2+^ concentration in the dense phase. We observed a substantially lower concentration of Mg^2+^ compared to the bulk concentration (**Fig. 2e**), suggesting that the condensed phase excluded Mg^2+^. These observations collectively confirm that the phase transition of aminoglycosides with RNA can encode Mg^2+^ concentration gradient, which modulates Mg^2+^ availability in the dilute phase, providing a fundamentally strategy for modulating Mg^2+^ signaling.

### Molecular dynamics simulation of aminoglycosides-RNA phase transition

To examine the competition of aminoglycosides and Mg^2+^ during RNA phase separation, we performed large-scale molecular dynamics simulations using the intermediate resolution model iConRNA with explicit Mg^2+^ ions^54^. This model can capture the balance of various interactions of RNA backbone and bases as well as dynamic local and long-range structural features in phase transition. In particular, iConRNA recapitulates the lower critical solution temperature phase separation of RNAs such as CAG repeats and its nontrivial dependence on sequence, length, concentration, and Mg^2+^ level^54^. We first developed compatible intermediate resolution models of aminoglycosides (see Methods, **Supplementary Fig. 2-5**), and then evaluated whether they could induce phase separation by interacting with RNA and how they compete with Mg^2+^ in the condensed phase. The simulations showed that, in the presence of both aminoglycoside molecules and Mg^2+^, rA200 RNA chains were efficiently bridged to form stable clusters (**Fig. 2f and Supplementary Fig. 6a**) with aminoglycoside molecules significantly enriched within the clusters. Density analysis reveals that Mg^2+^ is excluded from the central region, where the aminoglycoside is enriched within the clusters (**Fig. 2f and Supplementary Fig. 6b**). This aligns with the experimental observation that aminoglycosides drive the release of Mg^2+^ once they undergo phase transition with RNAs. Direct co-existence simulations further confirmed the exclusive effect from various aminoglycosides, showing that the extent of Mg^2+^ release depends on the charge state of the aminoglycosides (**Fig. 2g and Supplementary Fig. 6c**).

### Gentamicin drives phase separation with mRNA in bacteria

To determine whether aminoglycosides can drive condensate formation in bacteria, we treated exponentially growing *E. coli* with different concentrations of gentamicin near its minimal inhibitory concentration and monitored dose-dependent condensate formation based on differential interference contrast using confocal microscopy. By quantifying the percentage of cells showing puncta formation, we identified ∼10 μM as the gentamicin concentration threshold for triggering condensate formation in bacteria with over 60% of cells showing puncta formation, while less than 10% of cells showed puncta at ∼5 µM (**Fig. 3a**). To confirm that the formed punctas are mediated by gentamicin, we conducted the same experiment using 1% labeled fluorescent GTIF594 and observed dose-dependent formation of gentamicin condensate (**Fig. 3a**). To further evaluate whether RNA is the key molecule condensed with gentamicin, we co-incubated 1% labeled GTIF594, gentamicin and RNAselect dye, and found that gentamicin and RNAs are co-localized together (**Fig. 3b**), confirming the formation of gentamicin-RNA condensates in living bacteria.

**Figure 3.**
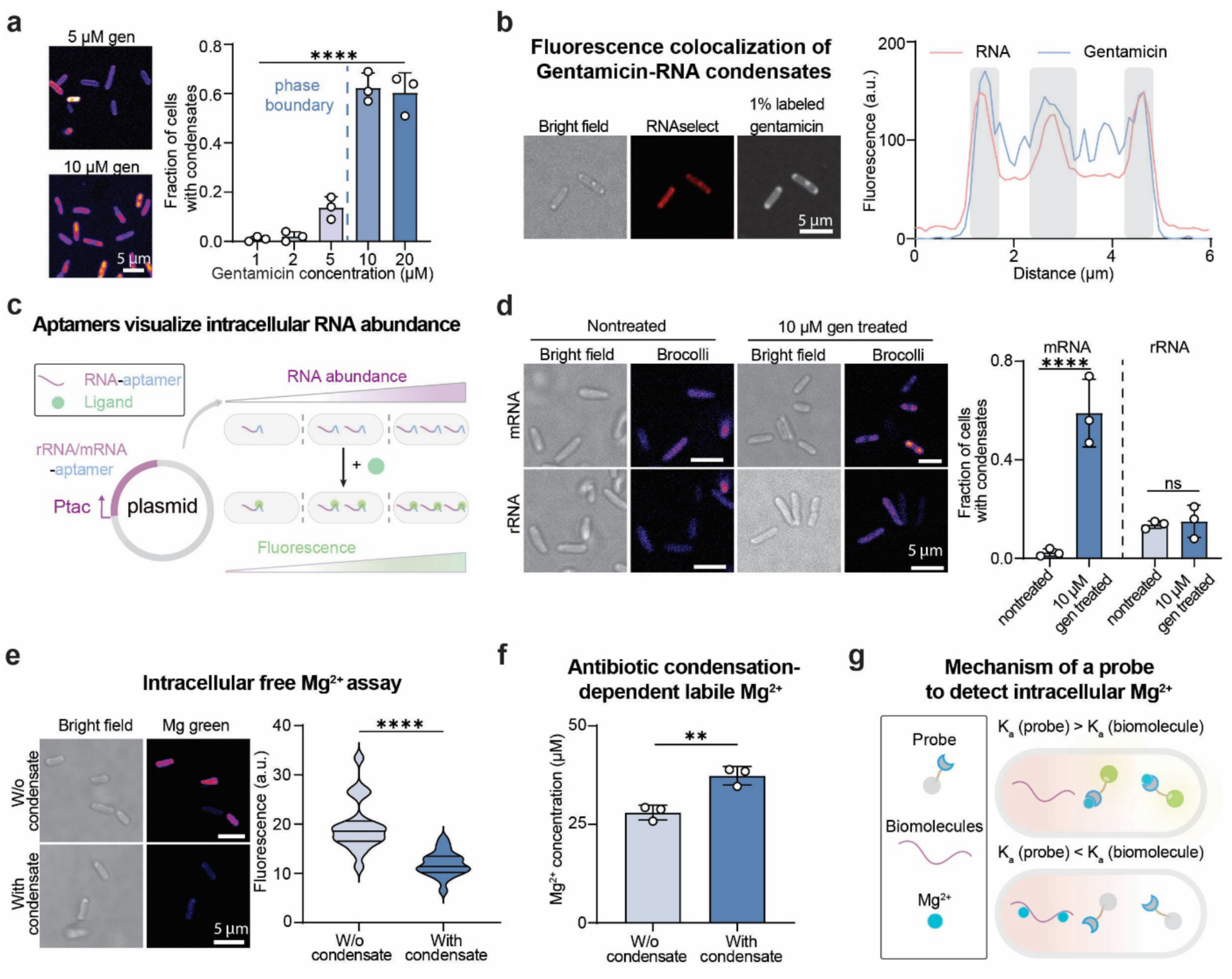
Antibiotics and mRNA drive condensate formation, modulating Mg^2+^ availability in *E. coli*. **a,** Evaluation of antibiotic–RNA condensate formation in cells following treatment with different concentrations of gentamicin. The left panel shows confocal fluorescence images of cells treated with 5 µM or 10 µM gentamicin. The right panel shows the fraction of cells containing condensates following treatment with different concentrations of gentamicin. More than 150 cells were quantified per biological replicate. **b,** Confocal fluorescence images showing the colocalization of gentamicin and RNA within gentamicin–RNA condensates in cells. The left panel displays the representative images showing gentamicin and RNA in the respective fluorescence channels. Scale bar represents 5 μm. The right panel shows the fluorescence intensity profiles as a function of distance, showing the spatial colocalization of gentamicin and RNA. Gentamicin and RNA are visualized using 1% (n/n) Texas Red-labelled gentamicin and 10 μM RNAselect, respectively. **c,** Schematics of the Broccoli RNA aptamer system for visualizing mRNA or rRNA levels. The Tac promoter controls transcription of the aptamer-tagged RNA, whose abundance is visualized upon addition of the fluorogenic ligand. Specific binding of the ligand to the Broccoli aptamer activates fluorescence, enabling RNA levels to be monitored. **d,** Visualization of intracellular mRNA or rRNA-aptamer distribution following the treatment of the gentamicin. The left panel displays the representative confocal images of mRNA and rRNA-aptamer distribution with or without treatment of gentamicin. Scale bar represents 5 μm. The right panel compares the fraction of cells containing the condensates with or without treatment of gentamicin. More than 100 cells were quantified per biological replicate. **e,** Evaluation of intracellular free Mg^2+^ concentration using Magnesium Green probe. The left panel displays the representative confocal fluorescence images of the cells with or without gentamicin-RNA condensates. The right panel compares the fluorescence of cells with or without condensates. More than 60 cells were quantified per biological replicate. **f,** Total cytoplasmic Mg^2+^ concentration of cells with or without antibiotic-RNA condensates. N = 3 biological replicates. Two-tailed t test for statistical analysis. **g,** Schematic of affinity-dependent competitive Mg^2+^ detection by a fluorescent probe. A high-affinity probe outcompetes biomolecules for target binding.

To evaluate the role of cellular RNAs in driving gentamicin condensate formation, we constructed a synthetic gene circuit that enables tunable control of RNA abundance based on the inducer concentration (**Fig. 3c**). To enable visualization of RNA distribution, we fused a small RNA aptamer, broccoli^55^, to a mCherry mRNA or rrlA rRNA lacking the ribosome-binding site. By inducing broccoli expression with distinct concentrations of inducers, we observed inducer concentration-dependent increase of total RNA concentration (**Supplementary Fig. 7a**). The addition of gentamicin converted the initially uniform distribution of mRNA into condensed structures, while rRNA remains diffusive (**Fig. 3d**). Consistently, testing other aminoglycosides showed that these antibiotics also induced mRNA condensation (**Supplementary Fig. 7b**). Together, these experiments support that mRNA serves as the key scaffold mediating aminoglycoside condensation in bacteria. This finding suggests that mRNA might serve as a responsive element to alter antibiotic efficacy.

### Phase transition of gentamicin induces Mg^2+^ signaling

We next investigated whether gentamicin-RNA condensates could modulate the spatial distribution of Mg^2+^ ions in bacteria. To this end, we first implemented magnesium green AM, which is a fluorogenic dye that binds to the labile Mg^2+^ ions^56^, to capture the free Mg^2+^ in cells treated with different concentrations of gentamicin. Interestingly, we found that the availability of labile Mg^2+^ decreased substantially once condensate formed (**Fig. 3e**). To evaluate whether the observed Mg^2+^ release through antibiotic condensation in vitro is valid in complex cellular matrix, we extracted cell lysate to evaluate whether gentamicin-RNA condensation could result in redistribution of Mg^2+^ using previously established protocol to separate the cytosolic environment and the condensed phase in bacteria lysate (**Supplementary Fig. 8**)^48^. This strategy releases biomolecule-bound Mg^2+^, allowing us to quantify the total asymmetric distribution of Mg^2+^ rather than only the labile pool. After condensate formation (at ∼10 μM gentamicin), we observed a noticeable increase in Mg^2+^ concentration in the cytosolic fraction (**Fig. 3f**). These two seemingly conflict observations suggest that the Mg^2+^ released during gentamicin–RNA condensate formation is subsequently exploited by other cellular machinery, rendering it inaccessible to detection by the competitive binding probe Magnesium Green AM (**Fig. 3g**). Together, both the experimental and computational studies confirm that aminoglycosides could regulate the distribution of Mg^2+^ as a strategy for electrochemical signaling^17,57^.

### Gentamicin condensates modulate ribosome activities

Since the gentamicin concentration needed to condense RNA in bacteria coincides with its minimal inhibitory concentration range for *E. coli* ^58^, we thus wondered whether this phase transition feature of gentamicin could be correlated with tolerance mechanism. We set out to investigate whether ribosome activity, which is known to be critical to drive antibiotic tolerance toward aminoglycosides and depends strongly on Mg^2+^ availability^27,59^, could be modulated by condensate-dependent ion signaling. To this end, we first used MG1655 strain with green fluorescent protein labeled onto the RplA, a core ribosomal subunit protein^60^ to analyze the spatial distribution of gentamicin and RplA in cells with gentamicin condensates, and found that though gentamicin segregate into a condensate but RplA remains relatively homogeneous (**Fig. 4a**). This observation suggests that though gentamicin condenses mRNAs, ribosome localization is largely unaffected by the condensation process. To evaluate whether the ribosome activity is modulated by antibiotic condensate-dependent signaling, we used the O-propargyl-puromycin based assay^61^ to analyze the ribosome activity by quantifying nascent protein synthesis rate. By comparing the translation rates in cells treated with different concentrations of gentamicin^48^, which create cells with or without condensates, we found that the translation rate closely mirrors the biphasic pattern of Mg^2+^ availability across the gentamicin concentration gradient and the cells with gentamicin condensates showed substantially increased translation rate (**Fig. 4b**). To further confirm this is Mg^2+^-dependent, we altered the amount of Mg^2+^ in the medium and observed the change of translation rate depending on the Mg^2+^ concentration under different conditions (**Supplementary Fig. 9**). These observations provide strong evidence to support our argument that phase transition of gentamicin activates Mg^2+^ signaling, which results in the enhancement of ribosome activity in a phase separation-dependent manner.

**Figure 4.**
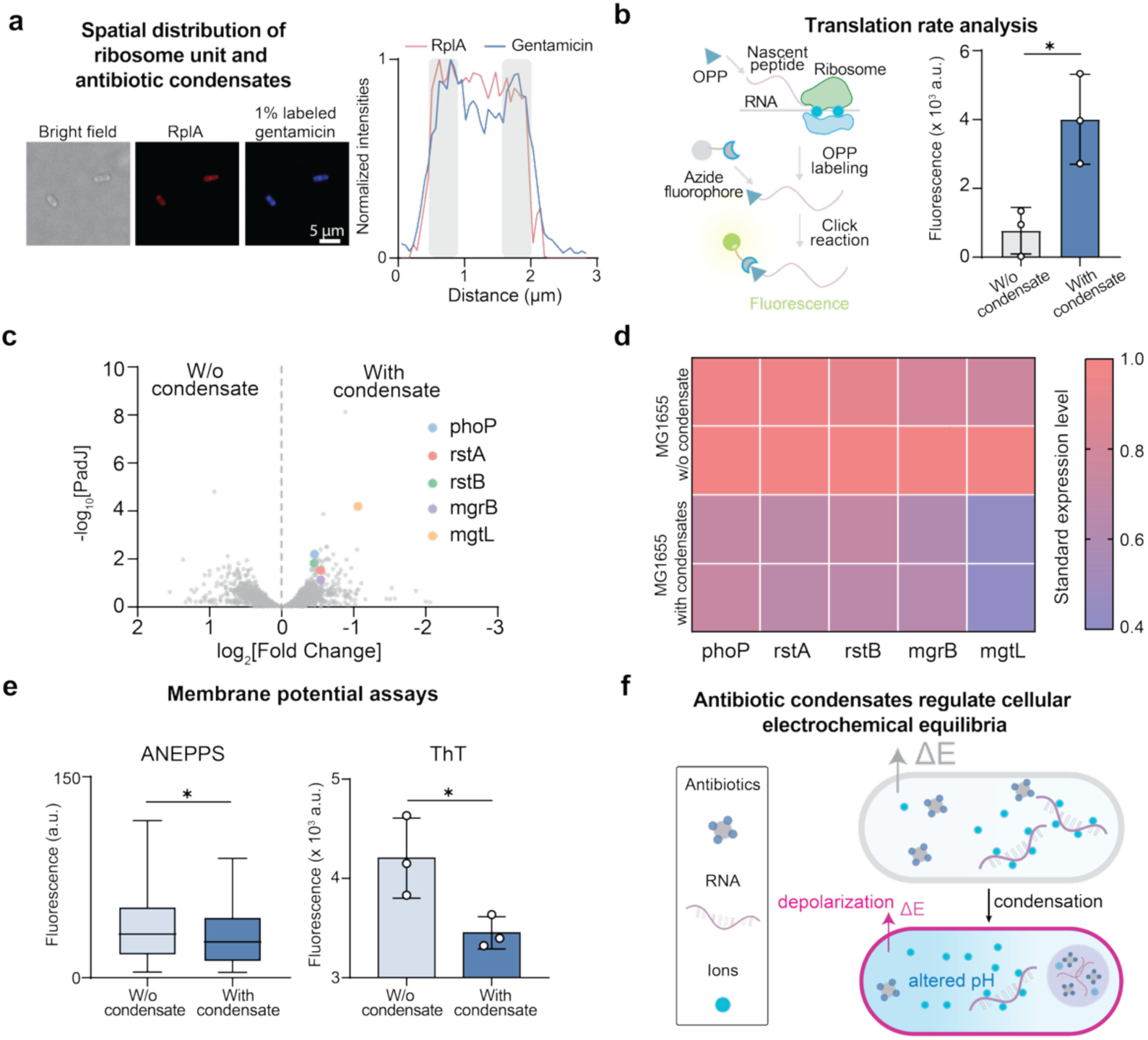
Antibiotic condensates regulate global cellular physiology by modulating ribosome activity, ion-dependent cellular processes and cellular electrochemical equilibria. **a,** Evaluation of the colocalization between ribosomes and gentamicin-RNA condensates. Left, representative confocal fluorescence images showing the intracellular distributions of gentamicin–RNA condensates and RplA, confirming non-colocalization between the two. Scale bar represents 5 μm. The right panel shows the fluorescence intensity profiles as a function of distance, showing the spatial non-colocalization of gentamicin-RNA condensates and RplA. **b,** Evaluation of translation rate of *E. coli* following the treatment of gentamicin. The left panel shows the illustration of O-propargyl-puromycin (OPP)-based measurement of intracellular protein synthesis. OPP is incorporated into nascent polypeptide chains during translation, and the incorporated OPP is subsequently labelled with a fluorescent probe through click chemistry. Fluorescence intensity reflects the amount of newly synthesized protein during the labelling period and is used as a measure of the cellular protein synthesis rate. The right panels compares the translation rate of *E. coli* with or without gentamicin-RNA condensates. Two-tailed t test for statistical analysis. **c,** Volcano plot (fold change of mRNA level between samples vs. adjusted p value) shows the distribution of transcriptomes in cells with and without gentamicin-RNA condensates. Statistically significant and featured genes are color coded based on the magnesium-homeostasis features. **d,** Relative transcript levels of magnesium-homeostasis genes in *E. coli* with or without gentamicin-RNA condensates. **e,** Evaluation of membrane potential of *E. coli* with or without gentamicin-RNA condensates. The ANEPPs assay is quantified using confocal microscopy. A higher fluorescence corresponds to membrane hyperpolarization. More than 80 cells were quantified per biological replicate. The ThT assay is quantified using flow cytometry. A higher fluorescence corresponds to membrane hyperpolarization. N = 3 biological replicates. Two-tailed t test for statistical analysis for both ANEPPs and ThT assays. **f,** Schematics of antibiotic-RNA condensation shifting intracellular labile ion contents, which mediates depolarization of membrane potential and modulates cellular pH condition.

### Gentamicin condensation-dependent activation of Mg^2+^ signaling

We next reasoned that if gentamicin condensates modulate Mg^2+^ signaling, their effects should extend beyond ribosome activity to other Mg^2+^-responsive cellular processes; investigating the Mg^2+^-dependent global cellular responses would distinguish a genuine Mg^2+^-mediated mechanism from gentamicin-specific ribosome action. To this end, we conducted RNA-seq analysis by comparing the expression profiles of cells treated with gentamicin at condition with or without condensates, at which gentamicin condensates appear at 10 µM but remain minimal at 5 µM. We observed that this transition was accompanied by a coordinated dampening of magnesium-starvation signaling. The PhoPQ two-component regulon^51,62^, which is the principal sensor of magnesium limitation in *E. coli*, was systematically lower in cells with condensates compared to those in cells without condensates. Specifically, the counts of *phoP* itself, its co-regulated targets *rstA* and *rstB*, and the PhoQ feedback regulator *mgrB* all decreased as condensates formed, with the regulon as a whole significantly shifted relative to the genome-wide background (**Fig. 4c and 4d**). The most pronounced change was *mgtL*, which is the key element reporting cytoplasmic Mg^2+^ status and gates expression of the *mgtA* importer^51,63^, which trended in the same direction (**Fig. 4d**). Because PhoPQ and *mgtA*/*mgtL* module are activated by magnesium scarcity, their coordinate repression upon condensate formation indicates that cells experience a more magnesium-replete cytoplasmic state. This transcriptional signature shows the downregulation of the magnesium-starvation alarm and deactivation of the induction of magnesium-import machinery by the presence of antibiotic condensates, which support our *in vitro* model that gentamicin-driven RNA coacervation liberates Mg^2+^ from RNAs.

### Gentamicin condensation alters cellular electrochemical equilibria

To further examine the potential effects of gentamicin-induced ion signaling, we next studied the changes of cellular electrochemical environments^48^. We reasoned that the formation of gentamicin-RNA condensates alters ion distribution in the intracellular space^48^, thus the global cellular electrochemical equilibria would be affected. To this end, we first evaluated the change of membrane potentials of cells under the with or without gentamicin condensates described previously. We employed two orthogonal fluorescence dye-based strategies: (i) the membrane-associated dye ANEPPS, which reports membrane hyperpolarization through fluorescence changes induced by the electronic Stark effect, and (ii) the positively charged dye ThT and negatively charged DiSBAC_2_(3), whose distribution is governed by the Nernst principle^20,48^. Both techniques reflect depolarization conditions when gentamicin condensate is formed (**Fig. 4e and Supplementary Fig. 10a-d**). This observation indicates that the release of Mg^2+^ mediates a more positive cellular interior, which depolarizes the membrane potential. This depolarized membrane potential suggests a change of environmental electrochemical driving force on modulating bacteria-chemical interactions^64,65^.

Next, we reasoned that protons and Mg^2+^ compete for the same anionic phosphate and carboxylate ligands^51^, so the two ions are linked through ion-exchange equilibria, and a redistribution of labile Mg^2+^ is expected to perturb proton equilibria on the negatively charged biomolecules in the cytosol ^66,67^, thereby mediating proton release (**Supplementary Fig. 10e**). To this end, we analyzed the intracellular pH environment using endogenously expressed ratiometric pH sensor, RpHluorin2^68^, whose emission wavelength depends on the proton availability in the cellular environment. Indeed, we found an increase of detectable proton availability in the cytosol upon condensate formation (**Supplementary Fig. 10f**). Together, this condensate-dependent change in proton dynamics further supports that condensate-dependent ion release can result in shift of cellular electrochemical environments (**Fig. 4f**).

### Gentamicin phase transition mediates multi-drug resistance

Since the biophysical, biochemical, and genetic changes induced by gentamicin condensation described above provide a mechanism for regulating cellular physiology during antibiotic treatment, we next investigated condensate-dependent antibiotic tolerance. To this end, we first tracked the growth recovery of cells treated with distinct gentamicin concentrations. We observed the gentamicin concentration-dependent non-monotonic, triphasic behaviors of recovery rate (**Fig. 5a and Supplementary Fig. 11a**). At gentamicin concentrations below the phase transition threshold, cellular growth was substantially reduced even though the antibiotic level remained below the minimal inhibitory concentration. In contrast, once the gentamicin concentration crossed the phase transition threshold, cellular regrowth became significantly enhanced, despite the antibiotic concentration being above the minimal inhibitory concentration. This observation directly aligns with condensate-dependent ribosome activity for enhancing antibiotic tolerance, demonstrating a novel tolerance mechanism that is independent of the presence of resistance gene^24^. Using antibiotics incapable of undergoing phase separation, we did not observe such non-monotonic growth effect (**Supplementary Fig. 11b**). Lastly, we quantified the probability of establishing tolerant behavior using 288 independently inoculated cultures treated with distinct gentamicin concentrations (**Supplementary Fig. 11c**). The same protocol was followed based on the seeding experimental design^69,70^. We found that more than 50% of cultures regrew at 10 μM gentamicin, a concentration at which condensates formed, whereas fewer than 1% regrew at 5 μM gentamicin. These observations suggest a critical role of antibiotic condensation on modulating drug tolerance.

**Figure 5.**
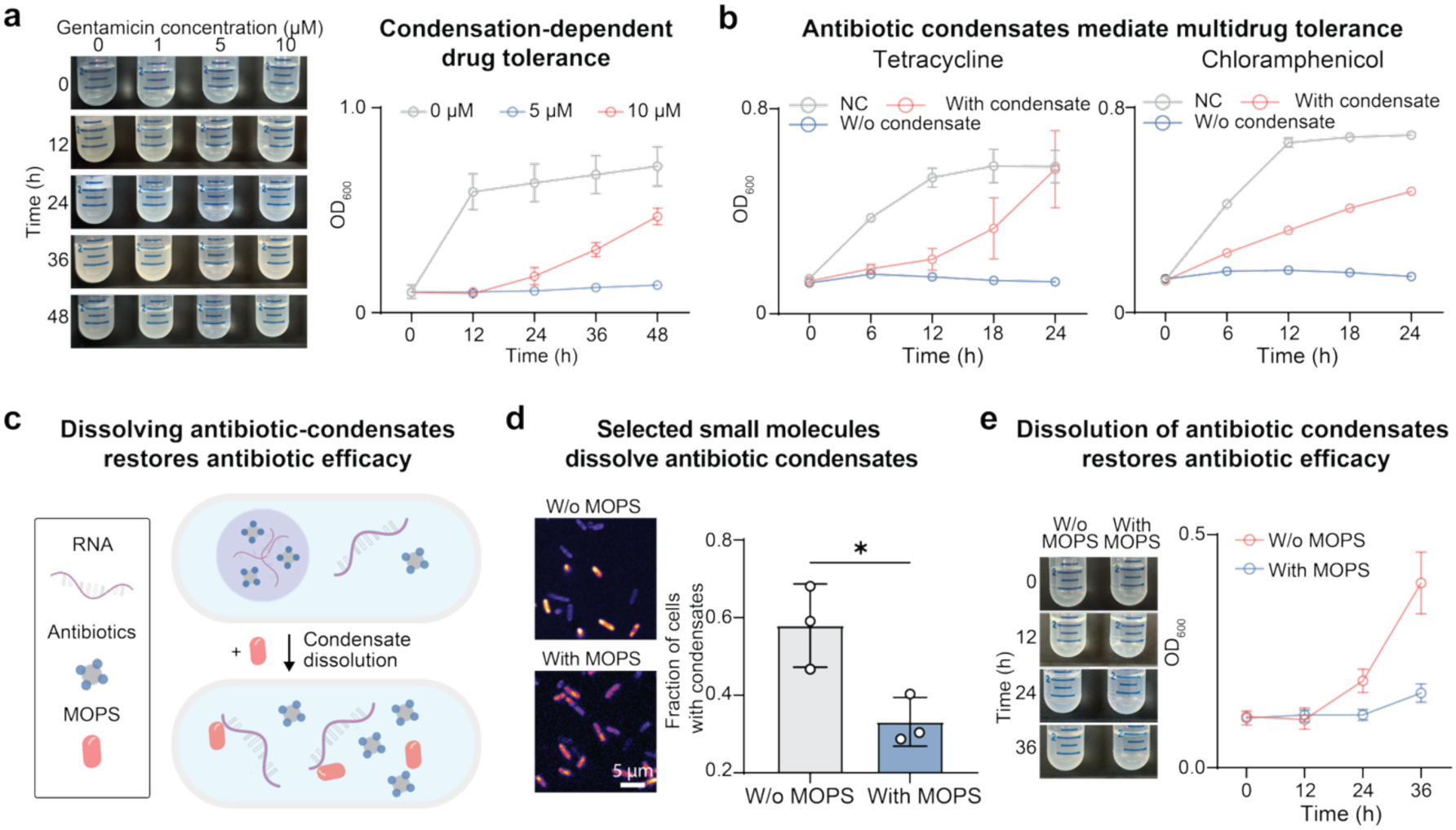
Antibiotic condensates encode non-monotonic multi-drug tolerance. **a,** Antibiotic-RNA condensates-dependent non-monotonic gentamicin tolerance. The left panel compares the growth conditions of *E. coli* treated with different concentrations of gentamicin. The right panel displays the corresponding growth curves of *E. coli* under indicated concentrations of gentamicin. Data is represented as mean ± SD. N = 3 biological replicates. **b,** Evaluation of multidrug tolerance of *E. coli* with or without gentamicin-RNA condensates. *E. coli* cells treated with 2 µm or 10 µM gentamicin represent condition without or with condensate formation, respectively. Cells are subsequently treated with sublethal concentration of tetracycline (2 mg/L) and chloramphenicol (5 mg/L). Data is represented as mean ± SD. N = 3 biological replicates. **c,** Schematics of drug disruption of antibiotic-RNA condensates. The drug is able to dissolve the condensates or prevent the formation of condensates by disrupting multivalent interactions between antibiotics and RNA. **d,** Evaluation of drug efficacy in disrupting gentamicin-RNA condensates. The left panel displays the representative fluorescence confocal images of cells treated with or without indicated drug. Scale bar represents 5 μm. The right panel quantifies the fraction of cells with gentamicin-RNA condensates. N = 3 biological replicates. Two-tailed t test for statistical analysis. **e,** Evaluation of the effect of drug treatment on the growth of *E. coli* treated with gentamicin. The left panel compares the growth conditions of *E. coli* treated with or without drug. The right panel displays the corresponding growth curves of *E. coli* under indicated conditions. Data is represented as mean ± SD. N = 3 biological replicates.

To confirm the role of ion signaling in mediating drug tolerance, we added Mg^2+^ into the medium and found that the recovery of cells was further enhanced (**Supplementary Fig. 11d**). Since this tolerance mechanism depends simply on the capability to drive labile Mg^2+^ ion-mediated ribosome activity, we reasoned that the release of free Mg^2+^ through phase separation might encode multidrug tolerance at the same time due to the crucial role of ribosome activity on mediating cell survival under antibiotic stress. We selected several ribosome-targeting antibiotics based on their distinct ribosome targeting sites, including tetracyclic and chloramphenicol, and co-treated *E. coli* with each of these antibiotics together with gentamicin. We leveraged the non-monotonic concentration dependence of cellular growth to analyze multidrug tolerance across different antibiotic concentrations (**Fig. 5b and Supplementary Fig. 11e**). Indeed, cells containing gentamicin condensates tolerated treatment with a second antibiotic despite being exposed to a higher total antibiotic dose.

### Disrupting condensates restore antibiotic efficacy

This condensate-dependent antibiotic tolerance mechanism suggests that perturbing condensates could potentially restore antibiotic efficacy. To test this concept, we added 3-Morpholinopropanesulfonic acid (MOPS), a zwitterionic buffer known to disrupt condensate formation in bacteria^71^, into the culture medium (**Fig. 5c**). We confirmed that the addition of MOPS did not impact cellular growth with growth kinetics comparable to control conditions (**Supplementary Fig. 11f**). In cells cultured in medium containing MOPS, at 10 µM gentamicin treatment, the percentage of cells with condensates drastically decreased (**Fig. 5d**), and their recovery was substantially delayed (**Fig. 5e and Supplementary Fig. 11g**). This experiment confirms a critical role of condensates on modulating drug tolerance. Similarly, when the condensates were dissolved by MOPS, the multidrug-tolerance phenotype observed under the same treatment conditions used in Fig. 5C was abolished. Together, these results suggest that targeting aminoglycoside–RNA condensates could provide a new strategy for modulating antibiotic tolerance and improving the treatment efficacy of combination antibiotic therapies.

## Discussions

Small molecules have long been thought to be organized primarily through reaction-dependent concentration gradients. We show that they can actively drive phase transitions that directly regulate their own spatial distributions and trigger downstream cellular functions. These results reveal phase transition as a previously underappreciated mechanism by which small molecules, which also possess multivalency through distinct functional groups, can encode cellular functions.

The condensate-driven ion release we describe offers a molecular explanation for a long-standing pharmacological paradox. Since the 1940s, numerous antibiotics have been reported to display a “paradoxical” or Eagle effect, in which bactericidal activity peaks at an intermediate concentration and then declines as the dose is raised further^72^. This non-monotonic dose–response has resisted a general mechanistic explanation, having been attributed variously to altered growth state, target saturation, or stress-response induction. Our results suggest that, for aminoglycosides, the paradox is an emergent property of a concentration threshold for phase separation: below the threshold the drug acts as a conventional translational inhibitor, whereas above it the same molecule co-condenses with RNA, liberating labile Mg^2+^ and sustaining ribosome activity even as free drug accumulates. This phenomenon also suggests a role of RNA in modulating antibiotic tolerance through modulating the spatiotemporal distribution of antibiotics through phase transitionThe triphasic growth-recovery profile we observed is the cellular signature of this transition, implying that the position of the survival window is set not by a resistance determinant but by the physicochemical phase behavior of the antibiotic itself. This finding also provides a resistance gene-independent mechanism by which antibiotic treatment can drive the de novo emergence of tolerant or resistant strains^69^.

These findings also reframe the relationship between an antibiotic’s concentration, its spatial organization, and its activity. The prevailing view treats small-molecule distribution as a passive consequence of reaction-diffusion gradients and electrochemical potentials. We instead find that an antibiotic can actively template its own compartmentalization through phase separation and, repurpose ions into signaling molecules. This places aminoglycosides within the electrochemical framework recently established for biomolecular condensates, which partition ions and establish interphase electric potentials and pH gradients that feed back on cellular physiology^45^. For example, the observed depolarization of membrane potential is expected to limit further proton-motive-force– dependent drug uptake^65^, so that crossing the phase boundary of antibiotic converts a lethal regime into a protective one. Thus, understanding how small molecules modulate cellular electrochemical equilibria provides a new dimension for understanding their functional roles in cellular physiology.

The mechanism we describe constitutes a route to multidrug tolerance that is entirely gene-independent. Classical resistance and most characterized tolerance mechanisms rely on dedicated genetic determinants, efflux pumps, modifying enzymes, target mutations, or persister-associated regulators^73^. In contrast, condensate-mediated Mg^2+^ release acts upstream of any single drug target: by sustaining ribosome activity and altering the cell’s electrochemical state, it protects against structurally distinct ribosome-targeting antibiotics co-administered with gentamicin. This carries two implications. Clinically, transient phenotypic tolerance can be elicited by the drug itself at concentrations near the minimal inhibitory concentration^74^, potentially contributing to treatment failure that genotype-based susceptibility testing would not predict, and cautioning that combination regimens built on ribosome-targeting agents could be mutually antagonized through this shared Mg^2+^/ribosome axis. Therapeutically, because the protective state depends on condensate formation, disrupting these condensates represents a generalizable strategy to restore efficacy. Finally, recognizing condensate-dependent electrochemical signaling as a component of antibiotic activity will establish a new theoretical framework for understanding both antibiotic mechanisms of action and the development of antibiotic tolerance in biology.

## Acknowledgements

Y.D. acknowledges the funding support from the National Institute of General Medical Sciences (grant number R35GM162028) and Alzheimer’s Association (grant number AARG-25-1486936). G.M.S. acknowledges the funding support from the National Institute of General Medical Sciences (grant number R35 GM139645), the Army Research Office (grant numbers W911NF261A170, W911NF-22-1-0107 and W911NF-1-0361) and the Bill and Melinda Gates Foundation (grant number INV-067331). J.C. acknowledges the funding support from the National Science Foundation (grant number CHE 25168941).

## Author contributions

Y.D. generated the idea. Y.D., G.M.S, and J.C. acquired the funding. Y.D., G.M.S, and Y.M. designed the project. Y.M., Y.D., G.M.S, J.C., and S.L. devised the experiment. Y.M., W.Y., E.M., Y.Z., J.W., J.C., Z.L., X.L., N.W., E.L. and E.B. performed the experiments. S.L. performed the simulation analysis. Y.M., J.W., Z.L., and E.L. analyzed the data. Y.D. and Y.M. wrote the manuscript. All authors read and/or edited the manuscript.

## Competing interests

All authors declare no competing interests.

## Data Availability

All data supporting the findings of this work are provided in the manuscript and its related Supplementary Information. Source data is provided for this paper.

## Notes

### Competing Interest Statement

The authors have declared no competing interest.

