## Supplementary Information for "Antibiotic condensates activate magnesium signaling to drive drug tolerance"

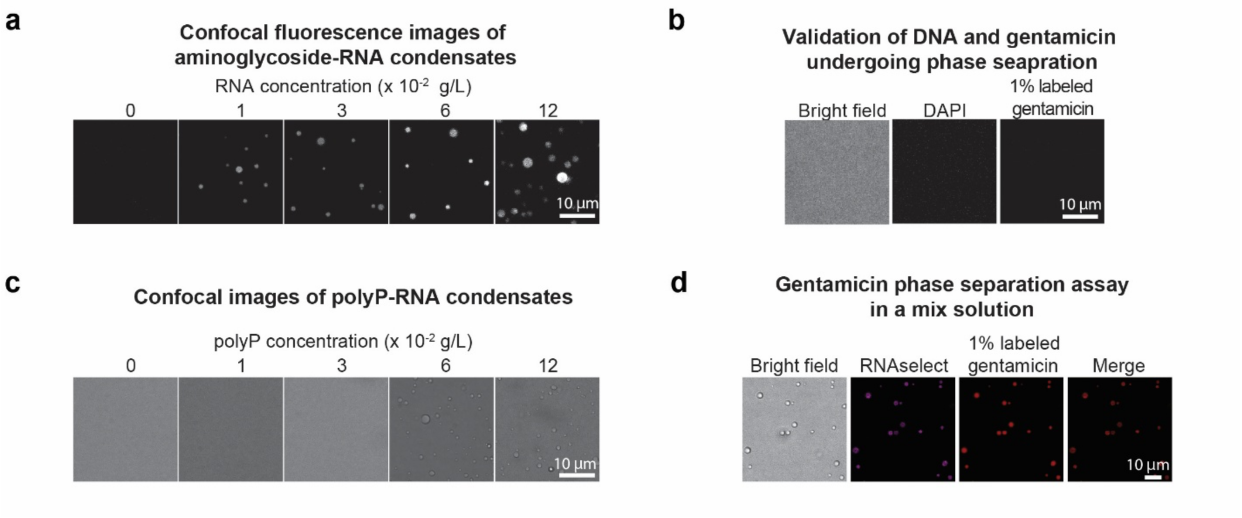


**Supplementary Figure 1. *In vitro* verification of antibiotic-RNA condensate formation.**

**a,** Confocal fluorescence images of *in vitro* gentamicin-RNA condensate formation at different concentrations of RNA and a fixed gentamicin concentration of 10 mM. Scale bar represents 10 µm.

**b,** Validation of gentamicin-DNA condensate formation. Confocal fluorescence images show that DNA and gentamicin are uncapable of undergoing phase separation *in vitro*. Gentamicin and DNA are visualized using 1% (w/w) Texas Red-labelled gentamicin and 0.5 µM DAPI, respectively. Scale bar represents 10 µm.

**c,** Validation of gentamicin-polyP condensate formation. Confocal images shows *in vitro* gentamicin-polyP condensates form only at a high concentration of polyP. Scale bar represents 10 µm.

**d,** Validation of gentamicin-RNA condensate formation in a DNA, RNA, and polyP mix solution. The final concentration of DNA, RNA, and polyP is 30 mg/L in each case. Confocal fluorescence images of gentamicin–RNA condensates shows the colocalization of gentamicin and RNA *in vitro*. Gentamicin and RNA are visualized using 1% (w/w) Texas Red-labelled gentamicin and 0.1 µM RNAselect, respectively. Scale bar represents 10 µm.

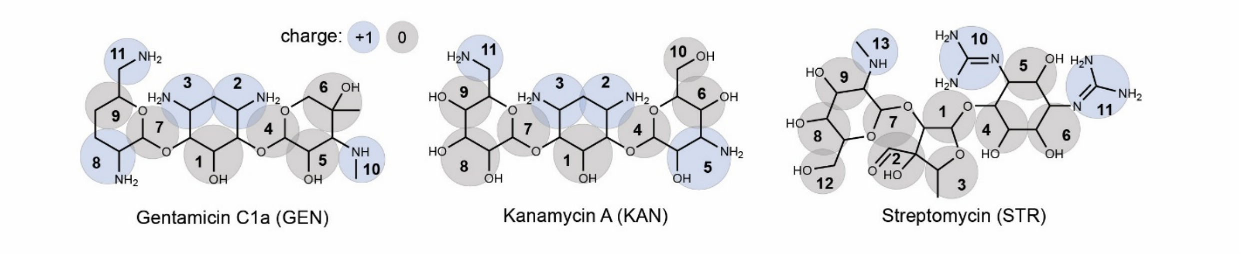


**Supplementary Figure 2. Chemical structures of the aminoglycoside antibiotics streptomycin, kanamycin, and gentamicin used in the simulation analysis.**

Groups highlighted in blue carry a single positive charge, while those highlighted in gray are uncharged.

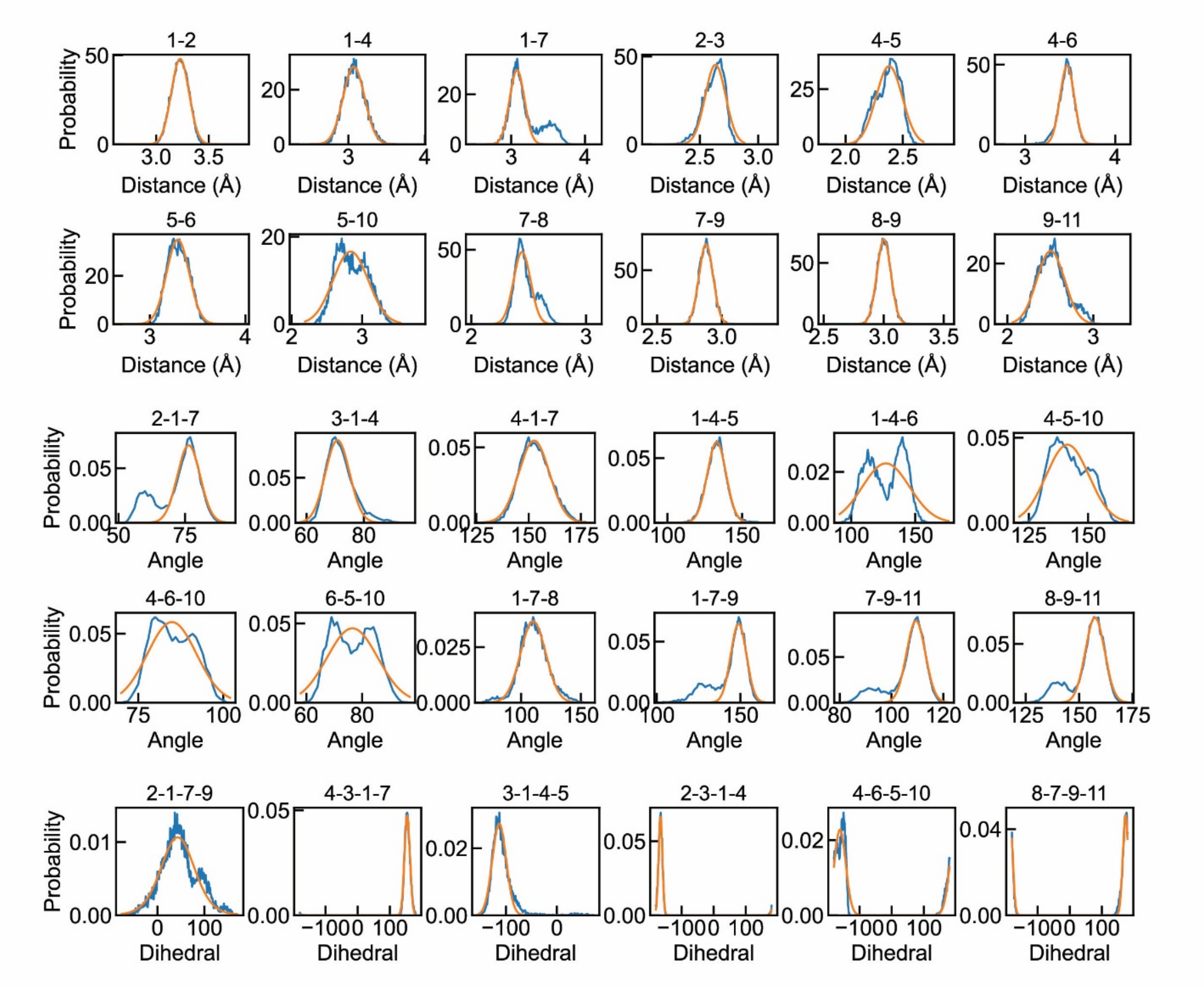


**Supplementary Figure 3.** **All-atom simulation-derived (blue) and fitted (orange) distributions of bonded terms for gentamicin C1a.**

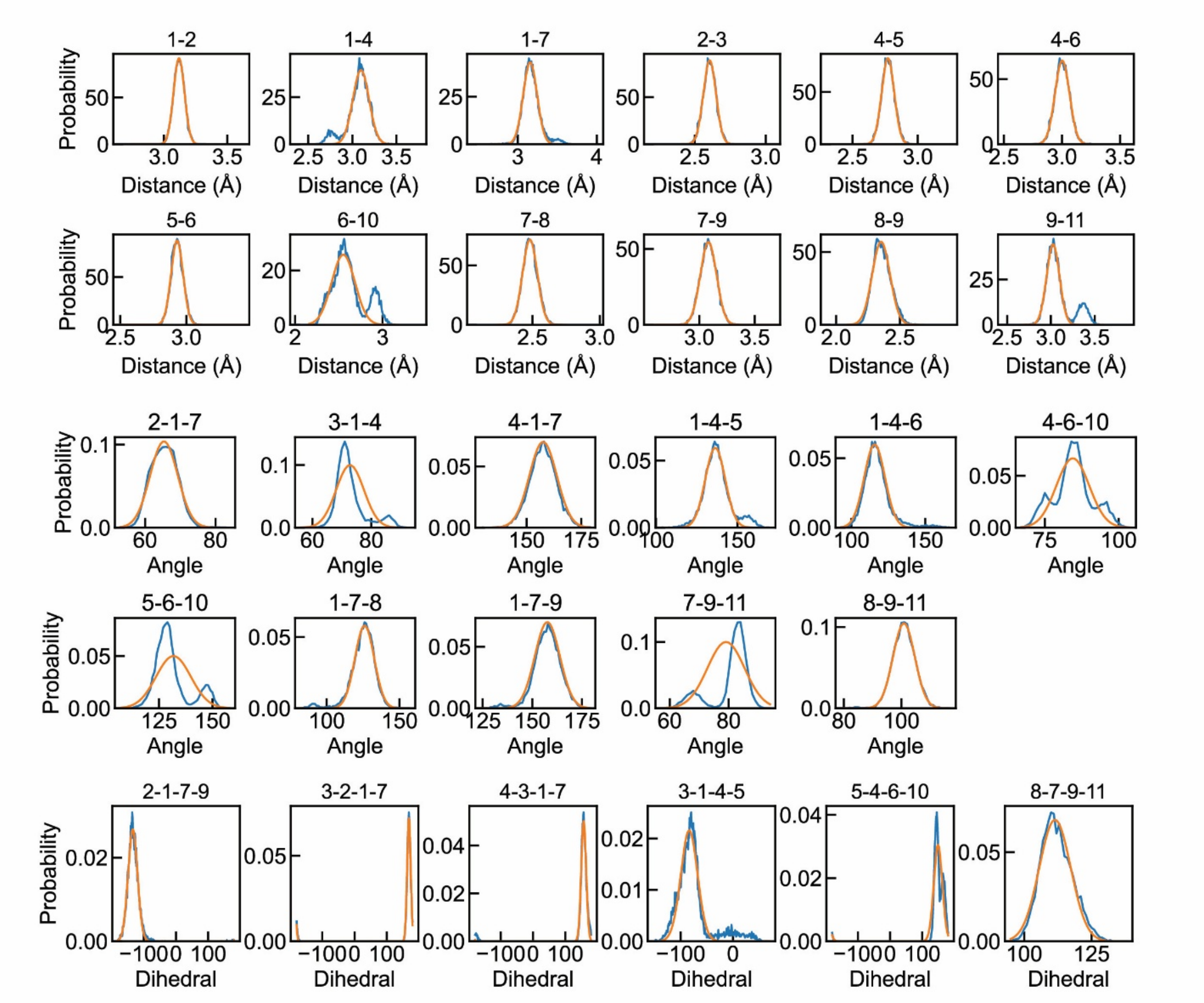


**Supplementary Figure 4. All-atom simulation-derived (blue) and fitted (orange) distributions of bonded terms for kanamycin A.**

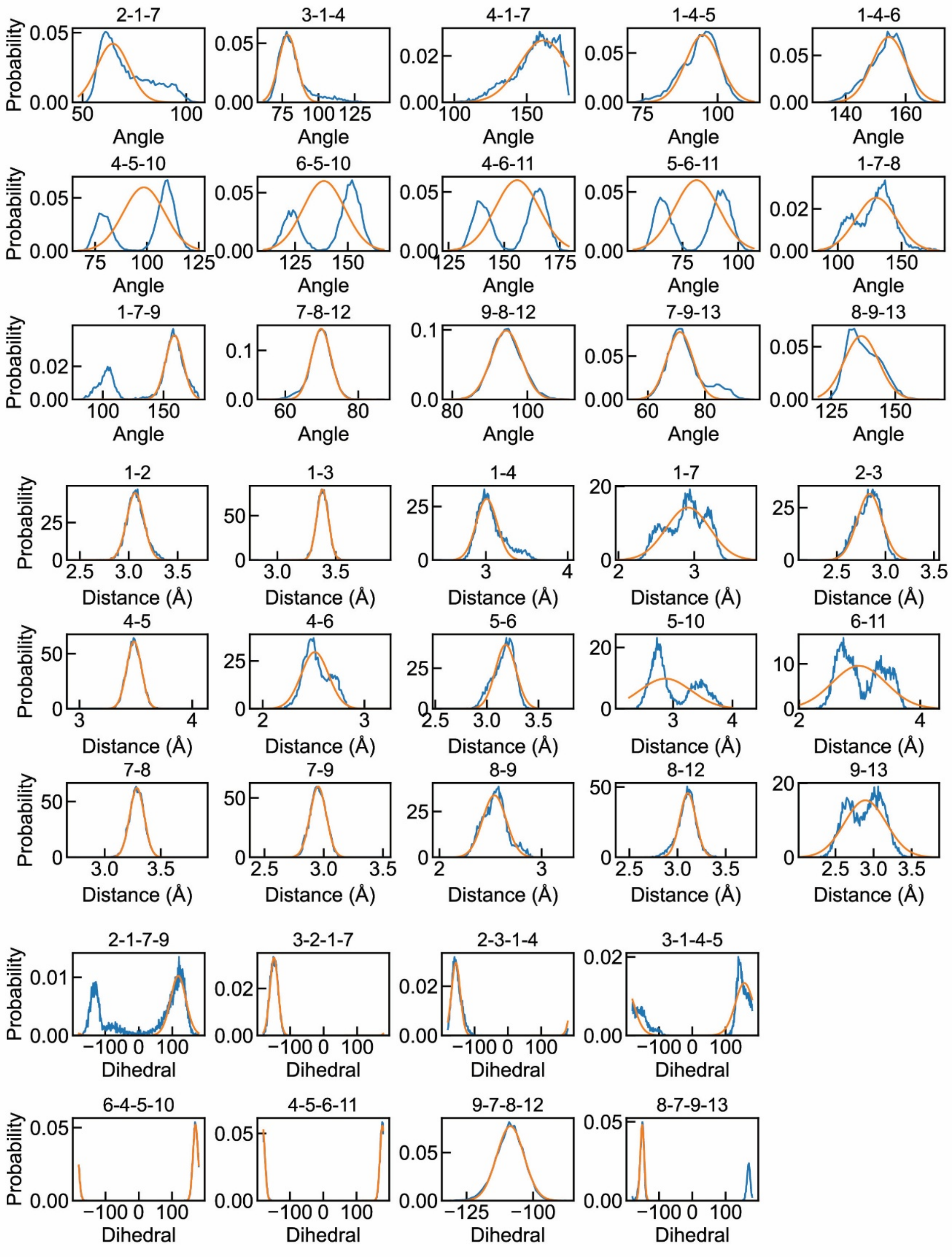


**Supplementary Figure 5.** All-atom simulation-derived (blue) and fitted (orange) distributions of bonded terms for streptomycin.

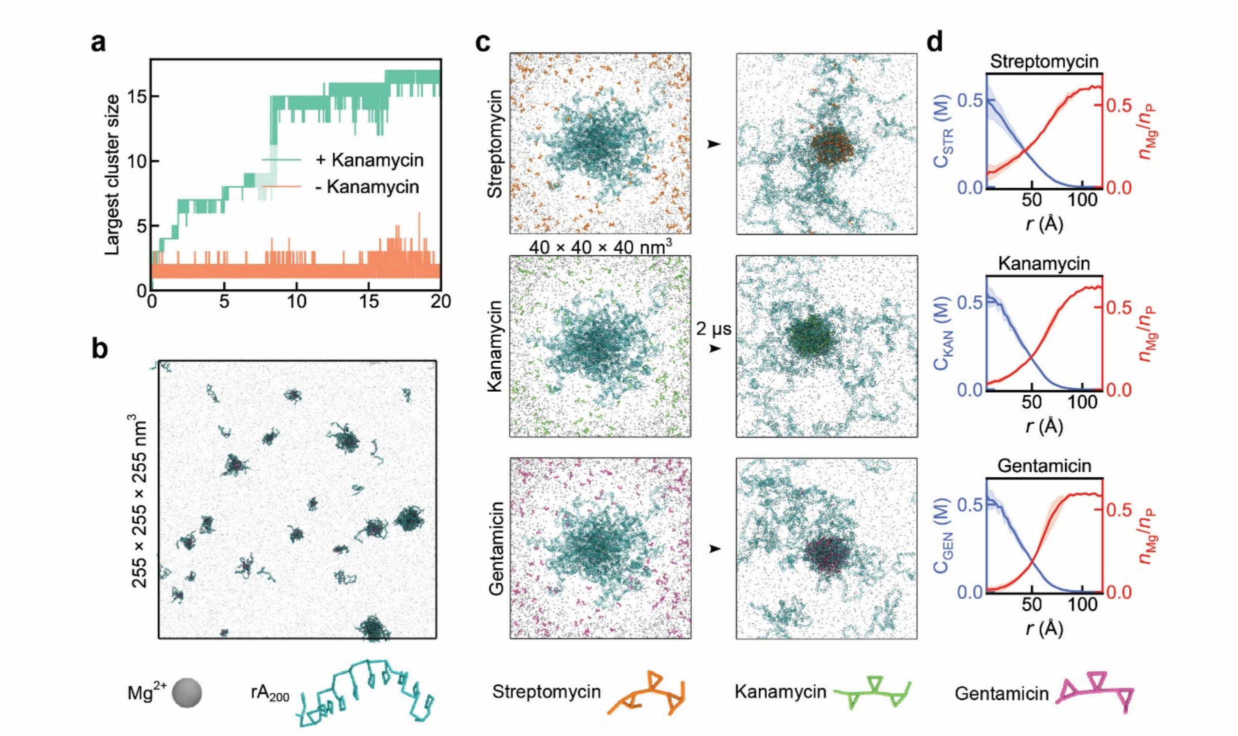


**Supplementary Figure 6. Simulation analysis of antibiotic-condensates.**

**a,** Kanamycin-induced RNA condensation kinetics over 20 μs simulation. This figure demonstrates the growth of the biggest cluster when kanamycin appears.

**b,** 20 μs simulation of 100 rA200 and 1000 kanamycin molecules.

**c,** 2 μs simulation of co-existence condensates using small cluster obtained in **Supplementary Fig. 6b** for kanamycin, gentamicin, and streptomycin, respectively.

**d,** Radial distributions of aminoglycoside concentrations (blue) and magnesium numbers per phosphate (red) for streptomycin, kanamycin, and gentamicin. Error bars (shadows) are estimated from block deviations.

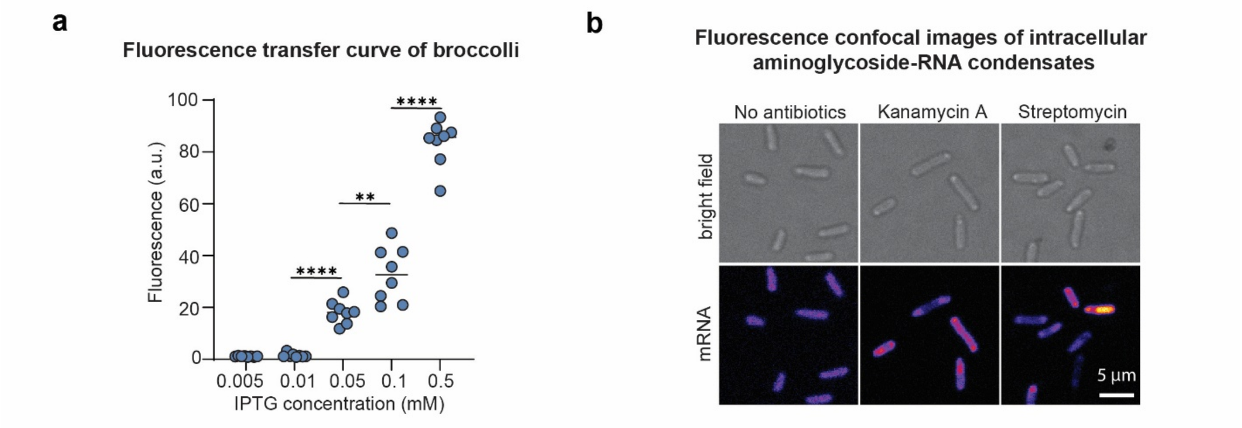


**Supplementary Figure 7. RNA aptamers enable visualization of intracellular RNA distribution.**

**a,** Evaluation of the transfer curve of the “Broccoli” expression system with P15A ori. N = 8 biological replicates. Two-tailed t test is used for statistical analysis.

**b,** Visualization of intracellular mRNA aptamer distribution following the treatment of the kanamycin A and streptomycin. Scale bar represents 5 µm.

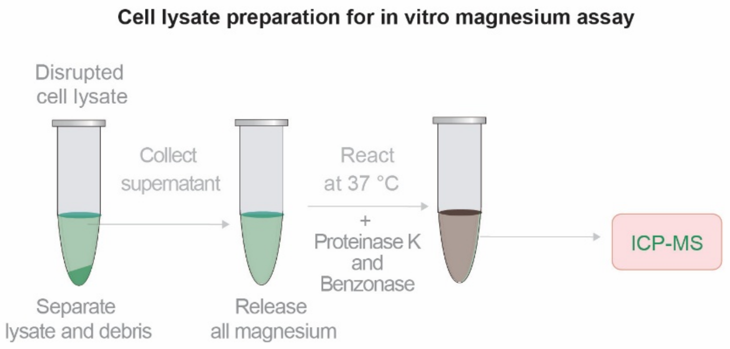


**Supplementary Figure 8. Workflow for preparing cell lysates for magnesium quantification.**

After 2 h of gentamicin treatment, cells are collected by centrifugation and resuspended in PBS to an $OD_{600}$ of 2. The 10 mL solution of cells are sonicated at 30% intensity using 1 s on/2 s off cycles. The lysates are then centrifuged to separate the supernatant and pellet fractions. The supernatant is treated with proteinase K and Benzonase and incubated at 37 °C for 1 h to release bound magnesium. Finally, the magnesium concentrations in all samples are measured using ICP–MS.

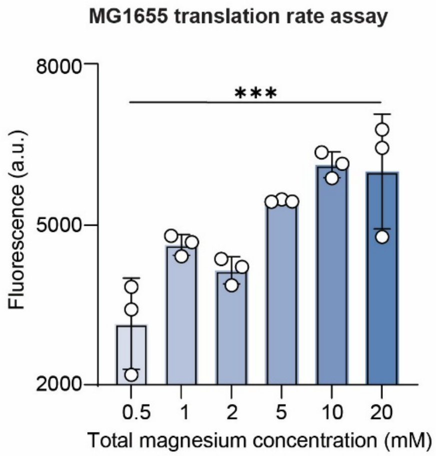


**Supplementary Figure 9. Validation of the positive correlation between magnesium concentration and intracellular translation rate.**

Single colonies of *E. coli* MG1655 are 1:100 into 4 mL of 2xYT medium supplemented with 0.4% glucose in 14 mL round bottom tubes and grown in a shaking incubator at 37 °C and 250 rpm for 12 h. Then, cultures are inoculated at a volume ratio of 1% (v/v) into the fresh M9 media until exponential phase ($OD_{600}$ = 0.25-0.3) is reached. Samples are removed from the cultures and diluted to an $OD_{600}$ of 0.1 in 1 mL of fresh M9 medium with indicated magnesium concentrations. Gentamicin is added to cell culture at a final concentration of 10 μM, and the antibiotic-treated cells are continually incubated at 37 °C for 1 h before collected for translation rate assay. Statistical significance was determined using an ordinary one-way ANOVA test.

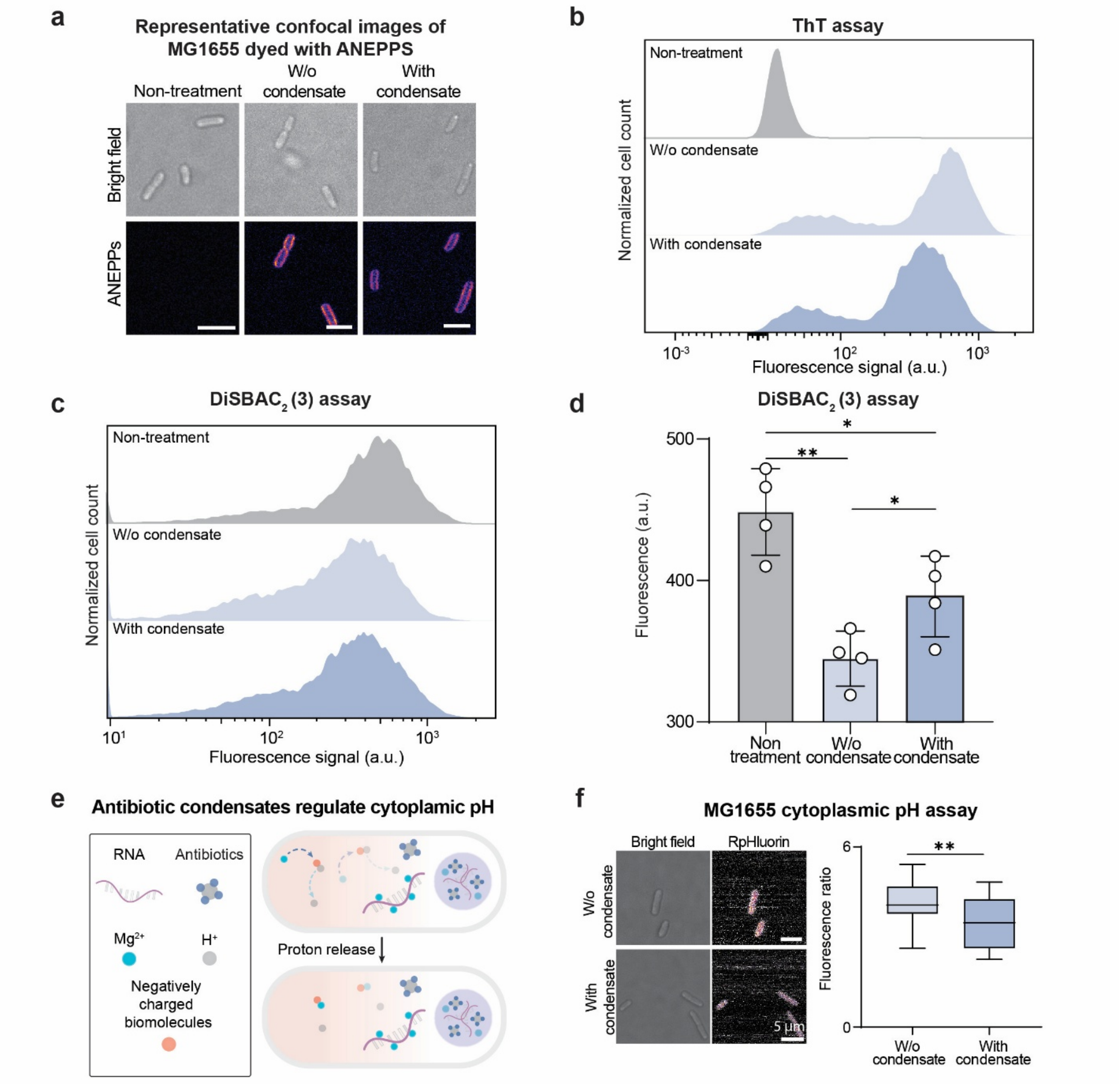


**Supplementary Figure 10. Physicochemical characterization of gentamicin-treated *E. coli* MG1655.**

**a,** Representative confocal fluorescence images showing the membrane potential of *E. coli* with or without gentamicin–RNA condensates using ANEPPS. Scale bar represents 5 μm.

**b,** Flow cytometry analysis of ThT (a cationic membrane potential dye) signal of single cells based on different conditions of MG1655 without or with gentamicin–RNA condensates. A higher fluorescence corresponds to membrane hyperpolarization.

**c,** Flow cytometry analysis of DISBAC$_2$(3) (an anionic membrane potential dye) signal of single cells based on different conditions of *E. coli* without or with gentamicin–RNA condensates. A higher fluorescence corresponds to membrane depolarization.

**d,** Evaluation of membrane potential of *E. coli* with or without gentamicin-RNA condensates using DISBAC$_2$(3). A higher fluorescence corresponds to membrane depolarization. N = 4 biological replicates. Two-tailed t test is used for statistical analysis for the DISBAC$_2$(3) assay.

**e,** Schematic illustration of the proposed mechanism by which antibiotic–RNA condensates regulate intracellular pH. The formation of these condensates displaces RNA-bound magnesium ions, promoting their release from RNA templates and altering intracellular magnesium homeostasis. This released magnesium may subsequently interact with other biomolecules to release hydrogen ion.

**f,** Evaluation of intracellular pH of *E. coli* with or without gentamicin-RNA condensates using RpHluorin2. The pH assay is quantified using confocal microscopy. A higher fluorescence ratio corresponds to an alkalic condition. Two-tailed t test is used for statistical analysis for the pH assay.

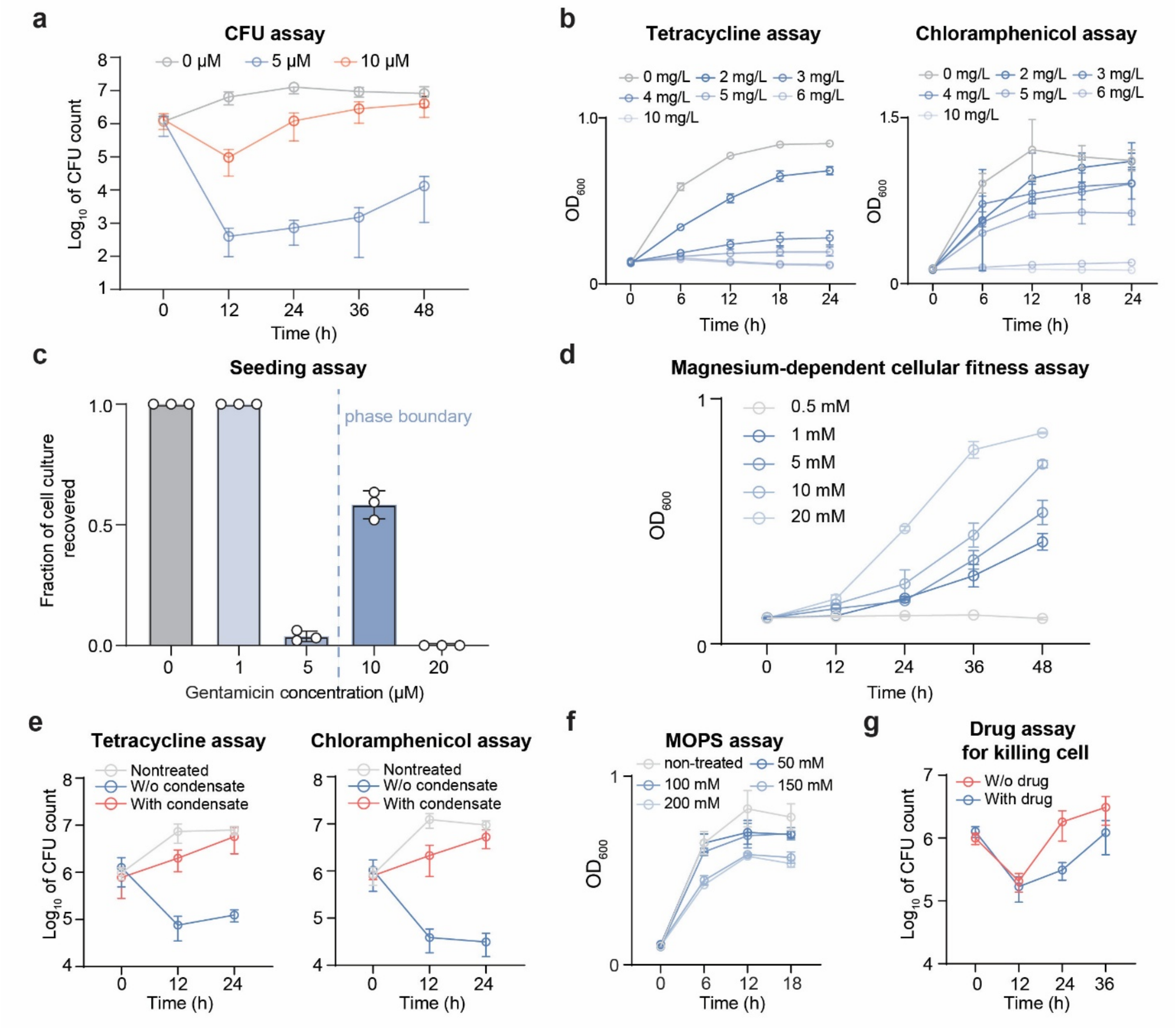


**Supplementary Figure 11. Evaluation of antibiotic-RNA condensates effects on cellular fitness.**

- **a,** Evaluation of Colony-forming units (CFUs) of gentamicin-treated *E. coli* at indicated time points. Data is represented as mean ± SD. N = 3 biological replicates.
- **b,** Evaluation of tolerance of *E. coli* to non-phase separable antibiotics. This figure compares the growth conditions of *E. coli* treated with different concentrations of tetracycline and chloramphenicol. Data is represented as mean ± SD. N = 3 biological replicates.
- **c,** Evaluation of condensate-dependent non-monotonic recovery. Different concentrations of gentamicin are added into *E. coli* cultures in 96-deepwell. After 48 h, cultures exhibiting visible turbidity are counted as positive for recovery. N = 288 replicates per condition.

**d,** Evaluation of effects of magnesium concentrations on gentamicin-treated *E. coli* growth. This figure compares the growth conditions of gentamicin-treated *E. coli* supplied with different concentrations of magnesium. Data is represented as mean ± SD. N = 3 biological replicates.

**e,** Evaluation of CFUs of tetracycline- and chloramphenicol-treated *E. coli* at indicated time points. Data is represented as mean ± SD. N = 3 biological replicates.

**f,** Evaluation of tolerance of *E. coli* to MOPS. This figure compares the growth conditions of *E. coli* treated with different concentrations of MOPS. Data is represented as mean ± SD. N = 3 biological replicates.

**g,** Evaluation of CFUs of gentamicin-treated *E. coli* with or without MOPS at indicated time points. Data is represented as mean ± SD. N = 3 biological replicates.

**Supplementary table 1. DNA sequences used in this study.**

| | |
|---|---|
| Broccoli | TTGCCATGTGTATGTGGGAGACGGTCGGGTCCATCTGAGACGGTCGGGTCCAGATATT<br>CGTATCTGTCGAGTAGAGTGTGGGCTCAGATGTCGAGTAGAGTGTGGGCTCCCACATA<br>CTCTGATGATCCAGACGGTCGGGTCCATCTGAGACGGTCGGGTCCAGATATTCGTATCT<br>GTCGAGTAGAGTGTGGGCTCAGATGTCGAGTAGAGTGTGGGCTGGATCATTCATGGCA<br>A |
| rrlA (rRNA) | GGTTAAGCGACTAAGCGTACACGGTGGATGCCCTGGCAGTCAGAGGCGATGAAGGA<br>CGTGCTAATCTGCGATAAGCGTCGGTAAGGTGATATGAACCGTTATAACCGGCGATTTC<br>CGAATGGGGAAACCCAGTGTGTTTCGACACACTATCATTAACTGAATCCATAGGTTAAT<br>GAGGCGAACCGGGGGAACTGAAACATCTAAGTACCCCGAGGAAAAGAAATCAACCG<br>AGATTCCCCCAGTAGCGGCGAGCGAACGGGGAGCAGCCCAGAGCCTGAATCAGTGT<br>GTGTGTTAGTGGAAGCGTCTGGAAAGGCGTGCGATACAGGGTGACAGCCCCGTACAC<br>AAAAATGCACATGCTGTGAGCTCGATGAGTAGGGCGGGACACGTGGTATCCTGTCTGA<br>ATATGGGGGGACCATCCTCCAAGGCTAAATACTCCTGACTGACCGATAGTGAACCAGT<br>ACCGTGAGGGAAAGGCGAAAAGAACCCCGGCGAGGGGAGTGAAAAAGAACCTGAA<br>ACCGTGTACGTACAAGCAGTGGGAGCACGCTTAGGCGTGTGACTGCGTACCTTTTGTA<br>TAATGGGTCAGCGACTTATATTCTGTAGCAAGGTTAACCGAATAGGGGAGCCGAAGG<br>GAAACCGAGTCTTAACTGGGCGTTAAGTTGCAGGGTATAGACCCGAAACCCGGTGAT<br>CTAGCCATGGGCAGGTTGAAGGTTGGGTAACACTAACTGGAGGACCGAACCGACTAA<br>TGTTGAAAAATTAGCGGATGACTTGTGGCTGGGGGTGAAAGGCCAATCAAACCGGGA<br>GATAGCTGGTTCTCCCCGAAAGCTATTTAGGTAGCGCCTCGTGAATTCATCTCCGGGG<br>GTAGAGCACTGTTTCGGCAAGGGGGTCATCCCGACTTACCAACCCGATGCAAACTGC<br>GAATACCGGAGAATGTTATCACGGGAGACACACGGCGGGTGCTAACGTCCGTCGTGA<br>AGAGGGAAACAACCCAGACCGCCAGCTAAGGTCCCAAAGTCATGGTTAAGTGGGAA<br>ACGATGTGGGAAGGCCCAGACAGCCAGGATGTTGGCTTAGAAGCAGCCATCATTTAA<br>AGAAAGCGTAATAGCTCACTGGTCGAGTCGGCCTGCGCGGAAGATGTAACGGGGCTA<br>AACCATGCACCGAAGCTGCGGCAGCGACACTATGTGTTGTTGGGTAGGGGAGCGTTC<br>TGTAAGCCTGTGAAGGTGTGCTGTGAGGCATGCTGGAGGTATCAGAAGTGCGAATGCT<br>GACATAAGTAACGATAAAGCGGGTGAAAAGCCCGCTCGCCGGAAGACCAAGGGTTC<br>CTGTCCAACGTTAATCGGGGCAGGGTGAGTCGACCCCTAAGGCGAGGCCGAAAGGC<br>GTAGTCGATGGGAAACAGGTTAATATTCCTGTACTTGGTGTTACTGCGAAGGGGGGAC<br>GGAGAAGGCTATGTTGGCCGGGCGACGGTTGTCCCGGTTTAAGCGTGTAGGCTGGTT<br>TTCCAGGCAAATCCGGAAAATCAAGGCTGAGGCGTGATGACGAGGCACTACGGTGCT<br>GAAGCAACAAATGCCCTGCTTCCAGGAAAAGCCTCTAAGCATCAGGTAACATCAAAT<br>CGTACCCCAAACCGACACAGGTGGTCAGGTAGAGAATACCAAGGCGCTTGAGAGAA<br>CTCGGGTGAAGGAACTAGGCAAAATGGTGCCGTAACTTCGGGAGAAGGCACGCTGA<br>TATGTAGGTGAAGCGACTTGCTCGTGGAGCTGAAATCAGTCGAAGATACCAGCTGGCT<br>GCAACTGTTTATTAAAAACACAGCACTGTGCAAACACGAAAGTGGACGTATACGGTGT<br>GACGCCTGCCCGGTGCCGGAAGGTTAATTGATGGGGTTAGCCGCAAGGCGAAGCTCT<br>TGATCGAAGCCCCGGTAAACGGCGGCCGTAACTATAACGGTCCTAAGGTAGCGAAAT<br>TCCTTGTCGGGTAAGTTCCGACCTGCACGAATGGCGTAATGATGGCCAGGCTGTCTCC<br>ACCCGAGACTCAGTGAAATTGAACTCGCTGTGAAGATGCAGTGTACCCGCGGCAAGA<br>CGGAAAGACCCCGTGAACCTTTACTATAGCTTGACACTGAACATTGAGCCTTGATGTG |

| | |
|---|---|
| | TAGGATAGGTGGGAGGCTTTGAAGTGTGGACGCCAGTCTGCATGGAGCCGACCTTGA<br>AATACCACCCTTTAATGTTTGATGTTCTAACGTTGACCCGTAATCCGGGTTGCGGACAG<br>TGTCTGGTGGGTAGTTTGACTGGGGCGGTCTCCTCCTAAAGAGTAACGGAGGAGCAC<br>GAAGGTTGGCTAATCCTGGTCGGACATCAGGAGGTTAGTGCAATGGCATAAGCCAGC<br>TTGACTGCGAGCGTGACGGCGCGAGCAGGTGCGAAAGCAGGTCATAGTGATCCGGT<br>GGTTCTGAATGGAAGGGCCATCGCTCAACGGATAAAAGGTACTCCGGGGATAACAGG<br>CTGATACCGCCCAAGAGTTCATATCGACGGCGGTGTTTGGCACCTCGATGTCGGCTCA<br>TCACATCCTGGGGCTGAAGTAGGTCCCAAGGGTATGGCTGTTCGCCATTTAAAGTGGT<br>ACGCGAGCTGGGTTTAGAACGTCGTGAGACAGTTCGGTCCCTATCTGCCGTGGGCGCT<br>GGAGAACTGAGGGGGGCTGCTCCTAGTACGAGAGGACCGGAGTGGACGCATCACTG<br>GTGTTCGGGTTGTCATGCCAATGGCACTGCCCGGTAGCTAAATGCGGAAGAGATAAGT<br>GCTGAAAGCATCTAAGCACGAAACTTGCCCCGAGATGAGTTCTCCCTGACTCCTTGAG<br>AGTCCTGAAGGAACGTTGAAGACGACGACGTTGATAGGCCGGGTGTGTAAGCGCAG<br>CGATGCGTTGAGCTAACCGGTACTAATGAACCGTGAGGCTTAACCTT |
| mCherry(mRNA) | ATGTCTGGTTCTCATCATCATCATCATCATGGTAGCAGCGGCGAAAACCTGTATTTTCAG<br>TCCATGGTGAGCAAGGGCGAGGAGGATAACATGGCCATCATCAAGGAGTTCATGCGC<br>TTCAAGGTGCACATGGAGGGCTCCGTGAACGGCCACGAGTTCGAGATCGAGGGCGA<br>GGGCGAGGGCCGCCCCTACGAGGGCACCCAGACCGCCAAGCTGAAGGTGACCAAG<br>GGTGGCCCCCTGCCCTTCGCCTGGGACATCCTGTCCCCTCAGTTCATGTACGGCTCCA<br>AGGCCTACGTGAAGCACCCCGCCGACATCCCCGACTACTTGAAGCTGTCCTTCCCCGA<br>GGGCTTCAAGTGGGAGCGCGTGATGAACTTCGAGGACGGCGGCGTGGTGACCGTGA<br>CCCAGGACTCCTCCCTGCAGGACGGCGAGTTCATCTACAAGGTGAAGCTGCGCGGCA<br>CCAACTTCCCCTCCGACGGCCCCGTAATGCAGAAGAAGACCATGGGCTGGGAGGCCT<br>CCTCCGAGCGGATGTACCCCGAGGACGGCGCCCTGAAGGGCGAGATCAAGCAGAGG<br>CTGAAGCTGAAGGACGGCGGCCACTACGACGCTGAGGTCAAGACCACCTACAAGGC<br>CAAGAAGCCCGTGCAGCTGCCCGGCGCCTACAACGTCAACATCAAGTTGGACATCAC<br>CTCCCACAACGAGGACTACACCATCGTGGAACAGTACGAACGCGCCGAGGGCCGCC<br>ACTCCACCGGCGGCATGGACGAGCTGTACAAGGGCAGC |
| RpHluorin2 | ATGGGCAGCAGCCATCATCATCATCATCACAGCAGCGGCCTGGTGCCGCGCGGCAGC<br>CATATGTCTAAAGGTGAAGAACTGTTCACCGGTGTTGTTCCGATCCTGGTTGAACTGGA<br>CGGTGACGTTAACGGTCACAAATTCTCTGTTTCTGGTGAAGGTGAAGGTGACGCTACC<br>TACGGTAAACTGACCCTGAAATTCATCTGCACCACCGGTAAACTGCCGGTTCCGTGGC<br>CGACCCTGGTTACCACCCTGTCTTACGGTGTTCAGTGCTTCTCTCGTTACCCGGACCAC<br>ATGAAACAGCACGACTTCTTCAAATCTGCTATGCCGGAAGGTTACGTTCAGGAACGTA<br>CCATCTTCTTCAAAGACGACGGTAACTACAAAACCCGTGCTGAAGTTAAATTCGAAGG<br>TGACACCCTGGTTAACCGTATCGAACTGAAAGGTATCGACTTCAAAGAAGACGGTAAC<br>ATCCTGGGTCACAAACTGGAATACAACTACAACGAACACCTGGTTTACATCATGGCTG<br>ACAAACAGAAAAACGGTACCAAAGCTATCTTCCAGGTTCACCACAACATCGAAGACG<br>GTGGTGTTCAGCTGGCTGACCACTACCAGCAGAACACCCCGATCGGTGACGGTCCGG<br>TTCTGCTGCCGGACAACCACTACCTGCACACCCAGTCTGCTCTGTCTAAAGACCCGAA<br>CGAAAAACGTGACCACATGGTTCTGCTGGAATTCGTTACCGCTGCTGGTATCACCCAC<br>GGTATGGACGAACTGTACAAACTCGAG |

**Supplementary table 2. Strains used in this study.**

| Strains | Genotype | Source |
|---|---|---|
| DH5α | *fhuA2::IS2 Δ(mmuP-mhpD)169 ΔphoA8 glnX44 ϕ80d[ΔlacZ58(M15)] rfbD1 gyrA96 luxS11 recA1 endA1 rphWT thiE1 hsdR17* | New England Biolabs |
| MG1655 | $F^{-}\lambda^{-} ilvG^{-}$ *rfb-50 rph-1* | American type culture collection |
| MG1655 (RplA-Gfp) | MG1655 *rplA::rplA-sfGFP* | Kind gift from Jacobs-Wagner lab |
